# Compartment-associated filtering and environmental gradients shape fungal communities in arid *Artemisia* species

**DOI:** 10.64898/2026.11.16.766971

**Authors:** Rezaul Karim, Rachid Elfermi, Ghita Amechatte, Fatima Zahra Aliyat, Mohamed Bammou, Mohamed Hijri, Bulbul Ahmed

## Abstract

Plant-associated fungal communities are shaped by interactions among host, environmental conditions, and compartment-specific filtering processes, yet the relative importance of these drivers remains poorly understood in arid and semi-arid ecosystems. We investigated fungal communities associated with three *Artemisia* species (*Artemisia herba-alba* Asso., *Artemisia mesatlantica* Maire and *Artemisia negrei* L.), including Moroccan endemics, across bulk soil, rhizosphere, root, and shoot compartments using ITS amplicon sequencing, culture-dependent isolation, and ecological analyses. Fungal community composition differed significantly among compartments (PERMANOVA, R² = 0.08, *P* = 0.012), whereas alpha diversity showed limited variation. Taxonomic profiles revealed compartment-associated shifts, with recurrent fungal genera concentrated in belowground compartments and a higher proportion of unclassified taxa in shoots. Core microbiome analyses identified overlapping fungal taxa between rhizosphere and root communities. Soil physicochemical gradients were associated with fungal community composition, and structural equation modeling indicated significant associations with both environmental variables and plant compartments. Source-tracking analysis attributed approximately 99.4% of root community composition to rhizosphere sources, whereas shoot communities showed contributions from both roots (60%) and the rhizosphere (39%). Culture-dependent isolation recovered 21 fungal isolates, with five genera shared between cultivation and sequencing datasets. Together, these findings reveal compartment-associated fungal community restructuring along the soil–plant continuum and highlight the complementary contributions of environmental conditions, plant-associated habitats, and cultivation-based characterization to understanding fungal microbiomes in arid ecosystems.

**Importance:** Understanding how fungal communities are organized across plant-associated habitats is essential for elucidating microbial assembly processes, particularly in arid ecosystems where environmental constraints may influence plant–microbe interactions. This study integrates ITS amplicon sequencing and culture-dependent approaches to investigate fungal communities associated with three *Artemisia* species, including Moroccan endemics. Our findings reveal significant compartment-associated differences in fungal community composition despite limited variation in alpha diversity. The recurrence of fungal taxa in rhizosphere and root compartments, together with source-tracking patterns, suggests progressive community restructuring along the soil–plant continuum. Soil physicochemical gradients were also associated with fungal community composition, highlighting the combined influence of environmental and plant-associated factors. Furthermore, the recovery of viable fungal isolates provides a complementary resource for future functional investigations. These findings extend our understanding of plant-associated fungal community organization in underexplored arid ecosystems and demonstrate the value of integrating cultivation and sequencing approaches in environmental microbiology.

## Introduction

Plant-associated microbial communities are essential for plant performance and ecosystem functioning, influencing nutrient acquisition, stress tolerance, and resistance to pathogen attacks (1). These microbiomes are not randomly distributed but are spatially and temporally structured across plant-associated compartments, including bulk soil, the rhizosphere, endophytic root tissues, and aerial organs (2). This spatial and temporal organization reflects the combined effects of environmental filtering, host selection and compartment-specific ecological constraints (3). While bacterial communities have been extensively characterized (4), fungal assemblages remain comparatively less understood despite their central roles in organic matter decomposition, nutrient cycling and plant-soil feedback, and plant health (5, 6). At broader biogeographical scales, this knowledge gap is even more pronounced in underexplored regions such as Africa, where soil fungal diversity remains poorly documented despite its ecological importance. A recent continent-wide survey of African soils revealed exceptionally high fungal diversity and strong spatial structuring across ecosystems, with diversity hotspots in savannas, temperate mixed forests, and dry tropical forests, and pronounced low-diversity zones in arid shrublands and deserts (7). That study further demonstrated that climatic variables such as precipitation and temperature, together with soil chemical properties, are key drivers of fungal diversity patterns across the continent, highlighting the strong influence of environmental filtering on large-scale fungal biogeography. Importantly, it also emphasized that African fungal communities remain significantly under-sampled relative to other continents, underscoring the urgency of expanding fungal biodiversity research to better understand, monitor, and conserve these essential microbial resources.

Building on these large-scale biogeographical patterns, it remains necessary to understand how such environmental filtering processes interact with host-associated selection mechanisms at finer ecological scales, particularly along the soil-plant continuum. A key ecological concept in plant microbiome research is the existence of a soil-plant continuum, where soil acts as a primary reservoir of microbial diversity and plant compartments impose progressively stronger selection filters. Within this continuum, the rhizosphere represents a first selective interface shaped by root exudation, while internal tissues impose further physiological and immunological constraints that strongly restrict microbial colonization. In parallel, soil physicochemical properties, particularly pH, nutrient availability, and texture, are well known to shape fungal community structure at broad spatial scales (3, 6, 8). However, the relative importance of abiotic environment gradients *versus* compartment-driven selection in structuring fungal communities along the soil–plant continuum remains insufficiently resolved, especially in natural and environmentally stressed ecosystems.

Arid and semi-arid ecosystems provide an ideal framework to address this gap due to strong environmental filtering, water limitation, and nutrient constraints that may intensify plant–microbe interactions. The genus *Artemisia* (Asteraceae), comprising over 500 species, is widely distributed across across arid and semi-arid regions and is known for its ecological resilience and adaptive strategies under harsh environmental conditions (9). In Morocco, *A. mesatlantica* Maire and *A. negeri* are endemic species, while *A. herba-alba* Asso. is characteristic components of steppe and dryland ecosystems, where they contribute to soil stabilization and ecosystem functioning (10, 11). Despite their ecological importance, the microbiomes associated with *Artemisia* species have been mainly studied in compartment-limited manner, often focusing on rhizosphere, and predominantly targeting bacterial communities (12). As a result, fungal community organization across the full soil-plant continuum remains poorly understood, particularly in endemic species and semi-arid environments. Here, we investigated fungal communities associated with three *Artemisia* species (*A. herba-alba*, *A. mesatlantica* and *A. negrei*) across bulk soil, rhizosphere, root and shoot compartments in an arid ecosystem of the High Atlas region of Morocco. Using fungal ITS amplicon sequencing, soil physicochemical characterization, and culture-dependent isolation, we characterized compartment-associated patterns of fungal community composition, identified core and indicator taxa, and quantified the relative contributions of environmental and compartment drivers to community assembly. We hypothesized that (i) fungal communities would exhibit strong compartmental structuring along the soil-plant continuum, (ii) rhizosphere and root compartments would contribute recurrent taxa reflecting selective recruitment, and (iii) soil physicochemical properties would contribute to community variation but be secondary to compartment effects, and (iv) culture-dependent and culture-independent approaches would recover overlapping but partially distinct fungal assemblages. This integrated framework provides new insights into the mechanisms governing fungal microbiome assembly in arid plant systems and highlights the dominant role of compartmental filtering in structuring plant-associated fungal communities.

## Materials and Methods

### Sample collection, processing, and soil physicochemical analysis

Bulk soil, rhizosphere, root, and shoot samples of three *Artemisia* species-*A. herba-alba* Asso. (widely distributed in Morocco), *A. mesatlantica* Maire (endemic to Morocco) and *A. negrei* L. (endemic to Morocco), were collected in June 2024 from the arid High Atlas region (latitude 31.85677–31.98795°N; longitude 4.99051–5.46546°W), including sites in Errachidia and Tinghir, Morocco. Plant identification was conducted using morphological characteristics and taxonomic keys, and subsequently verified by Professor Hamid Khamar (Scientific Institute, Rabat, Morocco). Voucher specimens were deposited in the National Herbarium of the Scientific Institute of Rabat under accession numbers RAB114680 (*A. herba-alba*), RAB114681 (*A. negrei*), and RAB114682 (*A. mesatlantica*). A total of 40 samples were collected, comprising four biological replicates per compartment (rhizosphere, root, and shoot) for each species, along with four bulk soil samples. Sampling was distributed across multiple sites, with *A. herba-alba* collected from two sites, *A. negrei* from one site, and *A. negrei* and *A. mesatlantica* co-occurring at a shared site. Samples were collected in sterile containers, transported on ice, and processed upon arrival.

Roots were gently washed under running water to remove adhering soil, rinsed with sterile distilled water, and air-dried under sterile conditions. Soil samples were sieved (2 mm mesh) to remove debris and plant material prior to physicochemical analysis. Soil properties were determined at the Agricultural Innovation and Technology Transfer Center (AITTC, Benguerir, Morocco), including pH, electrical conductivity (EC), soil texture (clay, silt, sand), total nitrogen (Total N), available phosphorus (P₂O₅), potassium (Total K), organic matter (OM), calcium carbonate (CaCO₃), nitrate (NO₃⁻), ammonium (NH₄⁺), chloride (Cl⁻), exchangeable oxides (K₂O, Na₂O, CaO, MgO), and trace elements (Cu, Mn, Fe, Zn, B). Analytical methods followed previously described protocols (4, 13).

### DNA extraction and ITS amplification

Plant tissues (root and shoot) were flash-frozen in liquid nitrogen and homogenized using a sterile mortar and pestle. Approximately 100 mg of homogenized material was transferred to sterile tubes and further disrupted using a TissueLyser II (QIAGEN, Germany). Genomic DNA was extracted using the DNeasy® Plant Pro Kit (QIAGEN, Germany) according to the manufacturer’s instructions. For bulk and rhizosphere soil samples, 250 mg of soil was processed using the DNeasy® PowerSoil® Pro Kit (QIAGEN, Germany). DNA quantity and purity were assessed using a BioSpectrometer (Eppendorf, Germany), and integrity was verified on 1% agarose gels. DNA extracts were stored at −20 °C until further processing. Fungal communities were amplified targeting the internal transcribed spacer (ITS) region using ITS-specific primers with CS adapters: ITS3 (TCGTCGGCAGCGTCAGATGTGTATAAGAGACAGAHCGATGAAGAACRYAG), and ITS4 (GTCTCGTGGGCTCGGAGATGTGTATAAGAGACAGTCCTCCGCTTATTGATAT GC). PCR reactions were performed in 25 μL volumes containing Platinum™ Direct PCR Universal Master Mix (Thermo Fisher Scientific, Morocco) and template DNA. Thermal cycling conditions consisted of an initial denaturation at 95 °C for 3 min, followed by 30–35 cycles of denaturation (95 °C, 30 s), annealing (50–55 °C, 30 s), and extension (72 °C, 45–60 s), with a final extension at 72 °C for 5 min. Amplifications were carried out using a Mastercycler X50s (Eppendorf, Germany). Amplicons were verified by electrophoresis on 1% agarose gels. Negative controls (non-template controls) were included in all PCR runs.

### Library preparation and amplicon sequencing

PCR products were purified using AMPure XP magnetic beads (Beckman Coulter, USA), washed with ethanol, and eluted in Tris buffer (10 mM, pH 8.5). A second PCR was performed to attach dual indices and Illumina sequencing adapters using indexing primers (Fluidigm, Canada). Indexed amplicons were purified, quantified using the Qubit dsDNA HS Assay Kit (Thermo Fisher Scientific), and pooled at equimolar concentrations. Sequencing was performed on an Illumina MiSeq platform (MegaFlex, Casablanca, Morocco) using the MiSeq Reagent Kit V3 (600-cycle paired-end reads).

### Sequence processing and statistical analysis

Raw sequencing data were processed using R (version 4.5.0). Base call files were converted to FASTQ format using the MiSeq Reporter pipeline. Sequence quality control and denoising were performed using the DADA2 package (14). Low-quality reads and sequences with expected errors greater than 2 were removed, and forward and reverse reads were truncated based on quality profiles. Chimeric sequences were identified and removed using the consensus method. Amplicon sequence variants (ASVs) were assigned taxonomic identities using the UNITE fungal ITS database (15, 16) with the assignTaxonomy function. Downstream analyses were conducted using the phyloseq package (17). Alpha diversity metrics (Shannon, Simpson and Observed) were calculated, and differences among compartments and species were assessed using analysis of variance (ANOVA) and non-parametric tests where appropriate. Beta diversity was calculated using Bray-Curtis dissimilarities and visualized using principal coordinates analysis (PCoA) with the vegan package (18). Differences in community composition were evaluated using PERMANOVA (ADONIS, 999 permutations). Relative taxonomic abundances were calculated using the *dplyr* package. Shared and unique taxa across compartments and species were visualized using Venn diagrams (*ggvenn* package). Associations between fungal communities and environmental variables were assessed using Mantel tests and visualized using *linkET*. Spearman correlation heatmaps were generated to evaluate relationships between taxa and soil properties. Core taxa were defined as ASVs with ≥70-90% prevalence and ≥0.1% relative abundance within a given compartment. Indicator species analysis was conducted using the *indicspecies* package based on the IndVal method (19). Differentially abundant taxa were identified using LEfSe implemented in the *microbiomeMarker* package (20). Structural equation modeling (SEM) was performed to evaluate relationships among soil properties, plant compartments, fungal diversity, and community composition, with model parameters estimated and evaluated based on standard goodness-of-fit criteria. Sankey diagrams illustrating taxon-specific transmission across compartments were generated using the *networkD3* package based on hierarchical Sankey representations (21). Statistical significance was determined at *p* < 0.05.

### Fungal isolation and Sanger sequencing

Fungal isolation from the rhizosphere, root and shoot samples of three *Artemisia* species was performed. Briefly, for epiphytic fungi, plant tissues (root and shoot) were rinsed three times with sterile distilled water to remove loosely attached debris, then were cut into ∼1 cm fragments. For the surface-associated fungal culture, the samples were directly plated onto Potato Dextrose Agar (PDA) medium under sterile conditions. For endophytic fungi, plant tissues were surface-sterilized using the immersion of samples in a 2% detergent solution for 3 minutes followed by rinsing with sterile water, and then 70% ethanol for 1 min approximately, followed by three successive rinses with sterile distilled water to remove residual ethanol. After sterilization, tissues were aseptically sectioned, the outer surface was removed, and internal fragments were directly placed onto PDA plates for endophyte isolation. For rhizosphere samples, approximately 1 g of soil was suspended in 9 mL of sterile saline solution and shaken at 150 rpm for 45 min to detach microbial cells. The suspension was serially diluted (10-¹-10-⁶), and pilot tests determined the optimal dilution for plating. Aliquots (100 μL) were spread onto PDA plates. Plates were incubated for 48-72 hours at 25°C. Distinct fungal culture were repeatedly recultured to obtain pure isolates. DNA was extracted using the DNeasy® PowerSoil® Pro Kit (QIAGEN, Germany) according to the manufacturer’s instructions. DNA quantity and purity were assessed using a BioSpectrometer (Eppendorf, Germany), and integrity was verified on 1% agarose gels. DNA extracts were stored at −20 °C until further processing.

For the amplification, ITS1 (5′-TCCGTAGGTGAACCTTGCGG-3′) and ITS4 (5′-TCCTCCGCTTATTGATATGC-3′) primer were used. PCR reactions (50 µL) contained 2 µL of each primer (10 µM), 8 µL GC enhancer reagent, 25 µL Master Mix ((Thermo Fisher Scientific, Morocco)), 1 µL of template DNA, and 12 µL nuclease-free water. Cycling conditions were as follows: 95°C for 5 min; 35 cycles of 95°C for 30 s, 57°C for 60 s, and 72°C for 60 s, followed by final extension at 72°C for 10 min. Amplifications were performed on a Mastercycler X50s (Eppendorf, Germany). Amplicons were verified on 1% agarose gels visualized with iBright FL1500 Imaging System (Thermo Fisher, Temara, Morocco). A non-template controls (NTC) were included in each run. Successful PCR products were purified and subjected to Sanger sequencing at Genome Quebec (Montréal, Canada). Forward and reverse sequences were assembled, quality-trimmed, and merged into contig using BioEdit (22). Taxonomic identification was performed using BLASTn against the NCBI nucleotide database.

### Phylogenetic analysis of fungal isolates

ITS consensus sequences of fungal isolates were compared against the NCBI nucleotide database using BLASTn. Closely related reference sequences were selected based on highest similarity, prioritizing curated and type-associated sequences when available. After removal of duplicates, the final dataset included 21 isolates and 116 reference sequences. Sequences were aligned using MUSCLE implemented in MEGA version 12 (23). Phylogenetic relationships were inferred using the maximum-likelihood method under the Tamura three-parameter model with gamma-distributed rate variation and a proportion of invariant sites (24). Branch support was assessed using 1,000 bootstrap replicates (25). Positions containing gaps or missing data were treated using partial deletion with a 95% site-coverage cutoff, resulting in a final alignment of 329 positions. The tree with the highest log-likelihood value was retained and visualized using the Interactive Tree of Life (iTOL) platform. Bootstrap values ≥70% were displayed, and study isolates were highlighted relative to reference sequences. The tree was interpreted as unrooted.

## Results

### Community composition differs among compartments in *Artemisia*

Alpha diversity did not differ significantly among compartments across *Artemisia* species-*Artemisia herba-alba* (Fig. S1), *Artemisia mesatlantica* (Fig. S2), and *Artemisia negrei* (Fig. S3). Most diversity indices (Shannon, Simpson, and Observed) showed non-significant differences (ANOVA/Kruskal-Wallis, *p* > 0.05), apart from Simpson index between rhizosphere-shoot and root-shoot in *A. herba-alba* (*p* = 0.0084). Despite the lack of statistical significance, a consistent decrease in richness and diversity from rhizosphere to shoot was observed across species. In contrast, beta diversity analysis revealed strong compartmental structuring of fungal communities (R^2^ = 0.08, F = 1.43, *p* = 0.012) (Fig. 1A). Principal coordinates analysis (PCoA) based on Bray-Curtis dissimilarities showed clear separation of rhizosphere, root and shoot samples along the first axis, which explained 12.9% of the variation, while the second axis accounted for 9.5%. In contrast, variation among *Artemisia* species was less pronounced, as indicated by overlapping distributions of species across compartments. Together, these results indicate that fungal diversity remains relatively stable across compartments, whereas community composition undergoes pronounced restructuring along the soil-plant continuum.

**Fig. 1.**
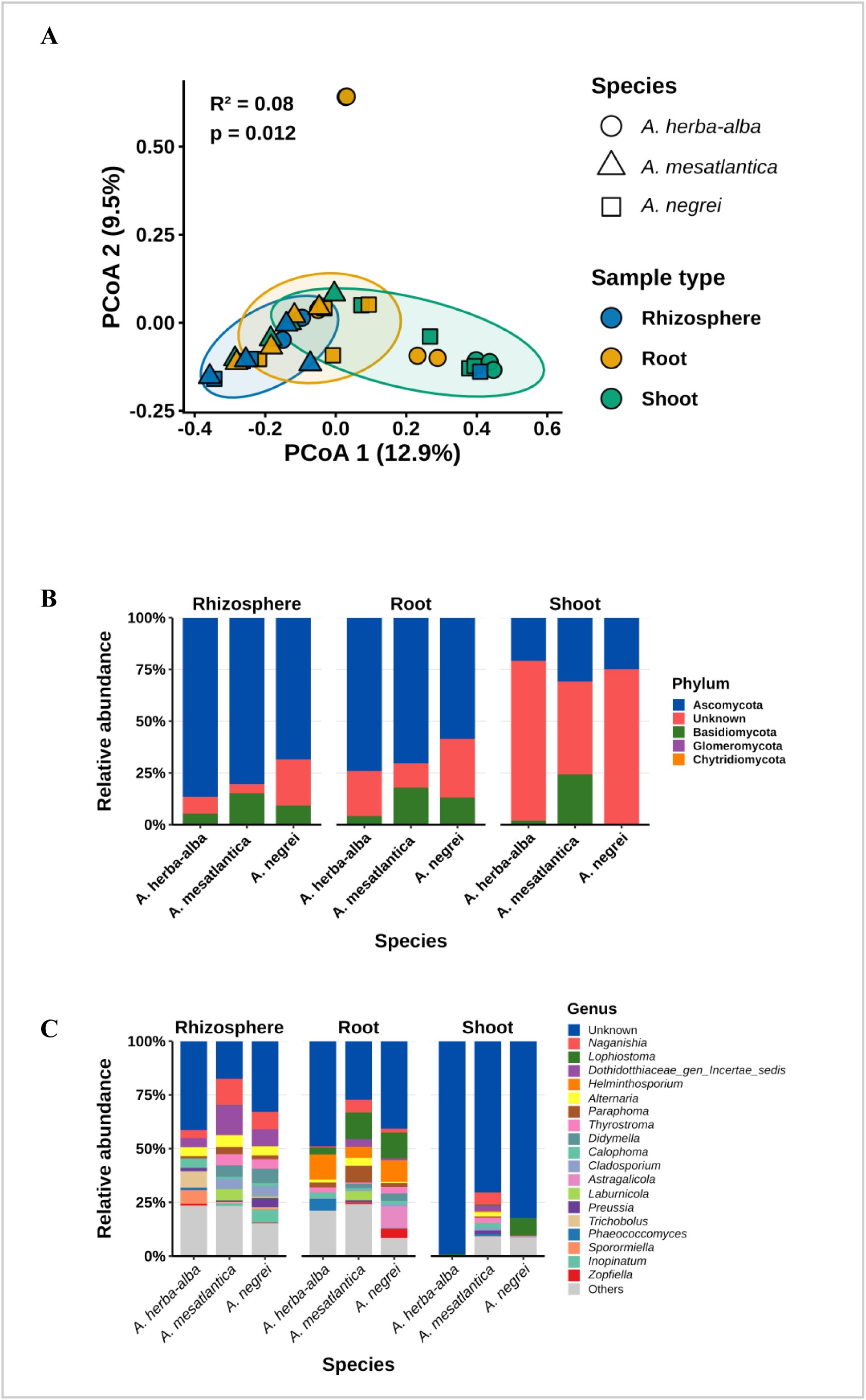
Fungal community structure and taxonomic composition across compartments. (A) Principal coordinates analysis (PCoA) of fungal community composition based on Bray-Curtis dissimilarities. (B) Relative abundance of major fungal phyla across rhizosphere, root, and shoot compartments. (C) Relative abundance of dominant fungal genera across compartments and *Artemisia* species.

### Taxonomic composition shifts along the soil-plant gradient

Fungal communities associated with *Artemisia* species were dominated by Ascomycota across rhizosphere and root compartments, whereas shoot communities showed increase in unclassified taxa and reduced representation of known fungal phyla (Fig. 1B). This pattern was particularly pronounced in *A. herba-alba* and *A. negrei*, where unclassified fungi accounted for most of the relative abundance in shoot samples. At genus level, communities were dominated by recurrent genera, including *Alternaria*, *Naganishia*, *Didymella*, *Lophiostoma* and *Cladosporium* (Fig. 1C). These taxa were detected across compartments but differed in relative abundance. Rhizosphere communities exhibited higher taxonomic heterogeneity compared to root communities, while genera such as *Paraphoma* and *Lophiostoma* were more abundant in root-associated samples. Shoot communities differed from belowground compartments by exhibiting a high proportion of sequences that could not be classified at the genus level and reduced representation of dominant genera. Notably, in *A. herba-alba* shoot samples, most sequences remained unclassified, indicating reduced taxonomic resolution in this compartment. Several taxa were abundant in rhizosphere, including members of *Calophoma*, *Laburnicola* and *Preussia*, showed reduced relative abundance in plant-associated compartments. These patterns indicate shifts in taxonomic composition along the soil-plant gradient.

### Rhizosphere and root compartments harbor consistent fungal taxa

Indicator species analysis identified several taxa associated with the rhizosphere across *Artemisia* species (Fig. S4A; Table S1). *Alternaria*, *Naganishia*, *Inopinatum*, *Spooneromyces* and members of Dothidotthiaceae (incertae sedis) showed high indicator values and were detected across host species, indicating strong compartment specific associations. No comparable indicator taxa were detected in other compartments, suggesting the weaker or less consistent associations outside the rhizosphere. In contrast, LEfSe analysis revealed broader patterns of differential enrichment across compartments (Fig. 2A) and host species (Fig. S4B). While rhizosphere communities were enriched in multiple taxa, additional taxa were also enriched in root and shoot compartments, reflecting more subtle differences in community composition. At host species level (Fig. S4B), distinct taxa were enriched in each *Artemisia* species, indicating secondary host-specific effects. Core microbiome analysis identified taxa consistently present within specific compartments (Fig. 2B). The core microbiome in the rhizosphere included *Alternaria*, *Naganishia*, *Didymella*, *Paraphoma*, *Cladosporium* and *Preussia*. In the root compartment, fewer taxa met the core threshold, with *Lophiostoma* representing the most abundant taxon, alongside *Naganishia*, *Alternaria* and *Paraphoma* (Table S2). Most root-associated core taxa were also present in the rhizosphere core microbiome, indicating substantial overlap between these compartments. These patterns indicate that both indicator and core taxa are concentrated in belowground compartments, with the strongest compartment specificity observed in the rhizosphere.

**Fig. 2.**
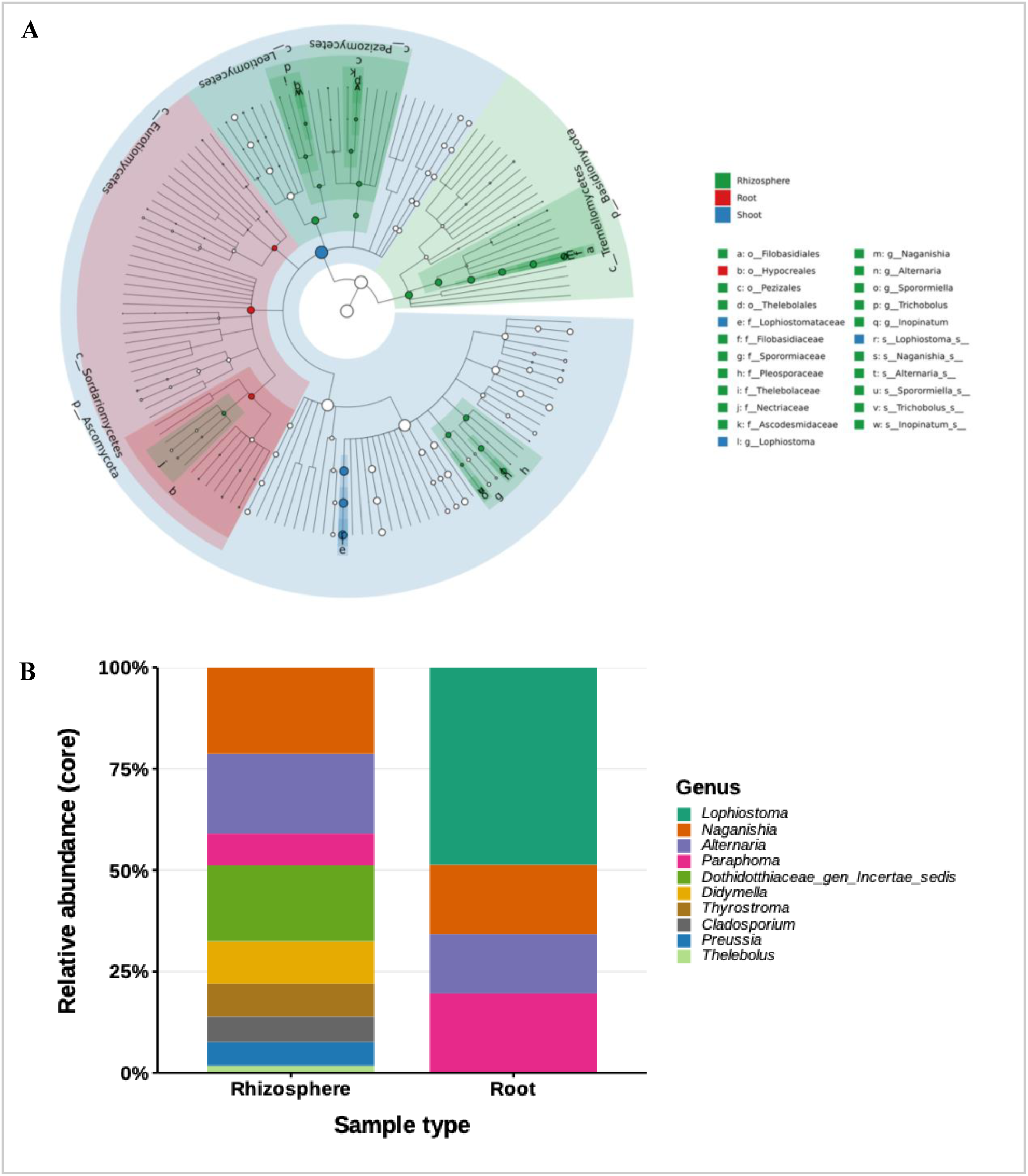
Compartment-associated fungal taxa. (A) LEfSe bar plot showing taxa significantly enriched in rhizosphere, root, and shoot compartments. (B) Core microbiome composition of rhizosphere and root compartments based on prevalence and relative abundance thresholds.

LEfSe analysis revealed differential enrichment patterns across plant compartments and host species (Fig. S4AB). At the compartment level (Fig. S4A), rhizosphere communities were enriched in multiple taxa, whereas root-associated enrichment was more restricted, and fewer taxa were enriched in shoots, consistent with lower diversity in this compartment. At the plant species level (Fig. S3B), distinct taxa were enriched in each *Artemisia* species, with *A. negrei* enriched in *Didymella, Thyrostroma*, and *Astragalicola*, *A. mesatlantica* in *Clonostachys* and *Dactylonectria*, and *A. herba-alba* in *Aureobasidium* and *Phaeoacremonium*. These results indicate that compartmental filtering is the primary driver of fungal community structure, with host species contributing to secondary differentiation.

### Fungal taxa and communities correlate with soil physicochemical gradients

Associations between fungal communities and soil physicochemical properties were assessed using Mantel tests, correlation analyses and dbRDA (Fig. 3; Fig. S5). Distance-based redundancy analysis indicated that the first two axes (dbRDA1 and dbRDA2) were associated with soil texture and pH-related variables. Species-specific patterns were observed, with *A. herba-alba* associated with pH, whereas *A. negrei* was associated with clay and silt fractions (Fig. S5). Mantel test analysis revealed significant correlations between fungal community composition and multiple soil physicochemical variables, with several variables showing strong associations (p < 0.01) (Fig. 3A), indicating that environmental gradients are linked to variation in community structure. Correlation analysis further revealed taxon-specific responses to environmental gradients (Fig. 3B). Genera such as *Didymella*, *Thyrostroma* and Dothidotthiaceae (incertae sedis) showed strong positive correlations with electrical conductivity, nitrate, carbonate content, organic matter, and multiple nutrient-related variables (e.g., total N, P₂O₅, K₂O, MgO, and micronutrients), while exhibiting negative correlations with pH. In contrast, *Apiosordaria* showed negative associations with most environmental variables. Other taxa, including *Alternaria*, *Calophoma*, *Naganishia*, *Lophiostoma*, *Leveillula*, and *Trichobolus*, displayed mixed patterns of association, whereas some genera exhibited weak or inconsistent relationships.

**Fig. 3.**
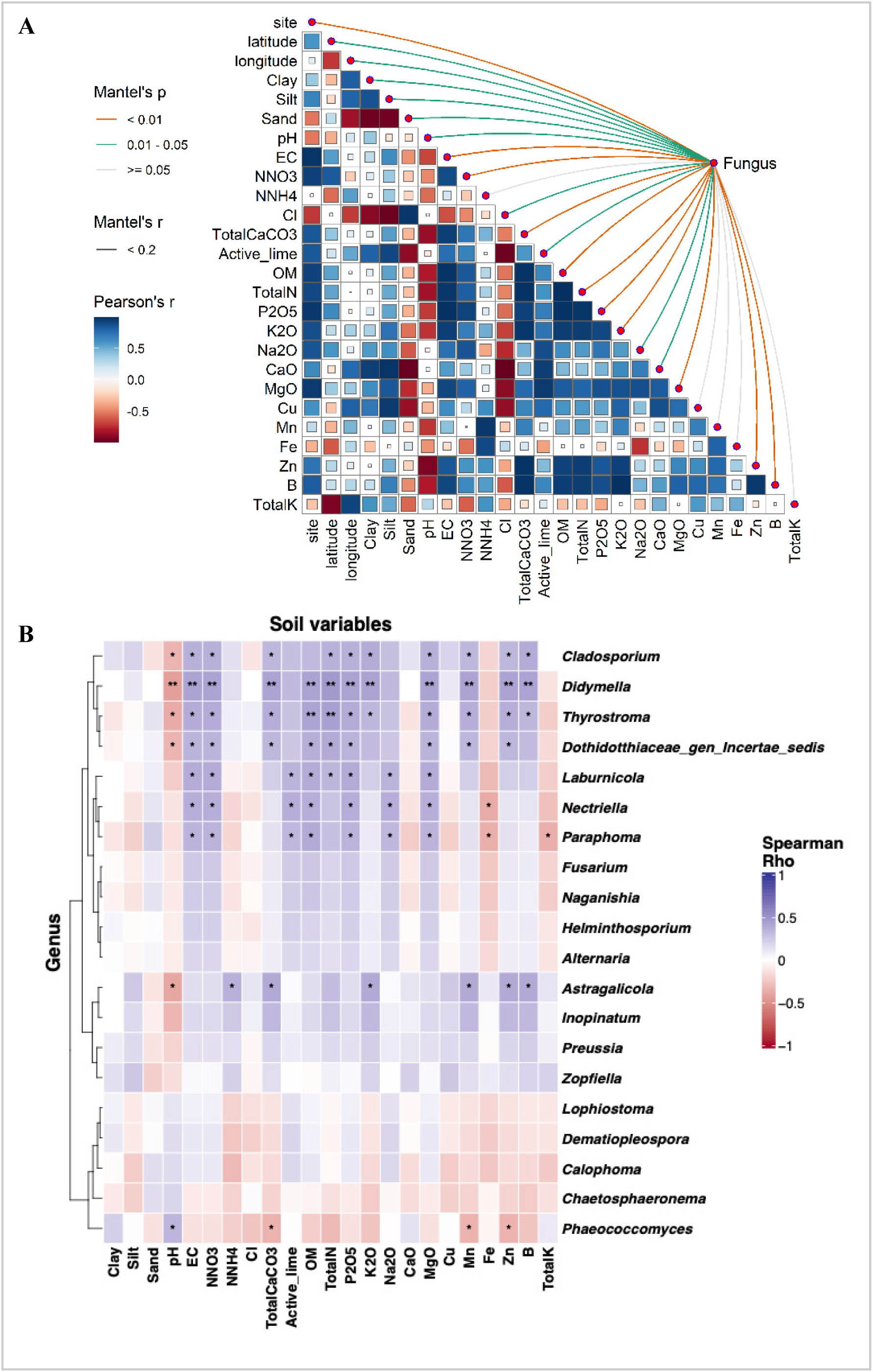
Relationships between fungal communities and environmental variables. (A) Mantel test showing associations between fungal community composition and soil physicochemical properties. (B) Spearman correlation heatmap of dominant fungal genera and environmental variables.

Comparison of fungal taxa among *Artemisia* species revealed both shared and species-specific components (Fig. S6A). A total of 37 taxa (23.1%) were common to all three species. Pairwise overlaps included 18 taxa (11.2%) between *A. negrei* and *A. mesatlantica*, 13 taxa (8.1%) between *A. herba-alba* and *A. mesatlantica*, and 6 taxa (3.8%) between *A. herba-alba* and *A. negrei*. Each species also contained unique taxa, with 32 taxa (20.0%) specific to *A. herba-alba*, 22 taxa (13.8%) to *A. negrei*, and 32 taxa (20.0%) to *A. mesatlantica*. These results indicate both a shared fungal core across species and a substantial proportion of host-associated taxa.

### Structural relationships among soil properties, compartments, and fungal communities

To further evaluate the relationships among environmental and compositional variables, structural equation modeling (SEM) was performed (Fig. S6B). Soil physicochemical properties, represented by principal components PC1 and PC2, were significantly associated with fungal community composition, with path coefficients of β = 0.037 (*p* = 0.031) for PC1 and β = −0.066 (*p* = 0.010) for PC2. In contrast, no significant association was observed between soil properties and fungal diversity (Shannon index). Plant compartment showed a significant relationship with community composition (*p* < 0.01), but not with fungal diversity. The model explained a greater proportion of variance in community composition (R² = 0.44) compared to fungal diversity (R² = 0.11). These results indicate that fungal community composition is associated with both environmental gradients and plant compartments, whereas fungal diversity shows limited responsiveness to these factors.

### Fungal communities follow a soil-rhizosphere-plant transmission pathway

Fungal communities showed a directional transition from rhizosphere to root and subsequently to shoot compartments (Fig. 4A). Root communities represented a subset of rhizosphere-associated taxa, whereas shoot communities exhibited further partitioning of these taxa, indicating progressive restructuring along the soil-plant continuum. Quantitative analysis further indicated that a relatively small fraction of taxa (∼12% of ASVs) was transmitted across compartments, yet these taxa accounted for more than half of the total abundance in shoot communities (∼51%). In contrast, a larger proportion of taxa was restricted to individual compartments but contributed minimally to overall abundance. Source contribution analysis further supported this pattern (Fig. 4B), indicating that root communities were primarily derived from the rhizosphere (∼99.4%), whereas shoot communities received contributions from both root (∼60%) and rhizosphere (∼39%) compartments. These results indicate that the rhizosphere acts as a major reservoir for downstream compartments. Species-specific analyses further showed similar transmission patterns across *Artemisia* species (Fig. S7A-C), where a small fraction of transmitted taxa contributed disproportionately to community abundance, indicating consistency in community transitions despite differences in host identity.

**Fig. 4.**
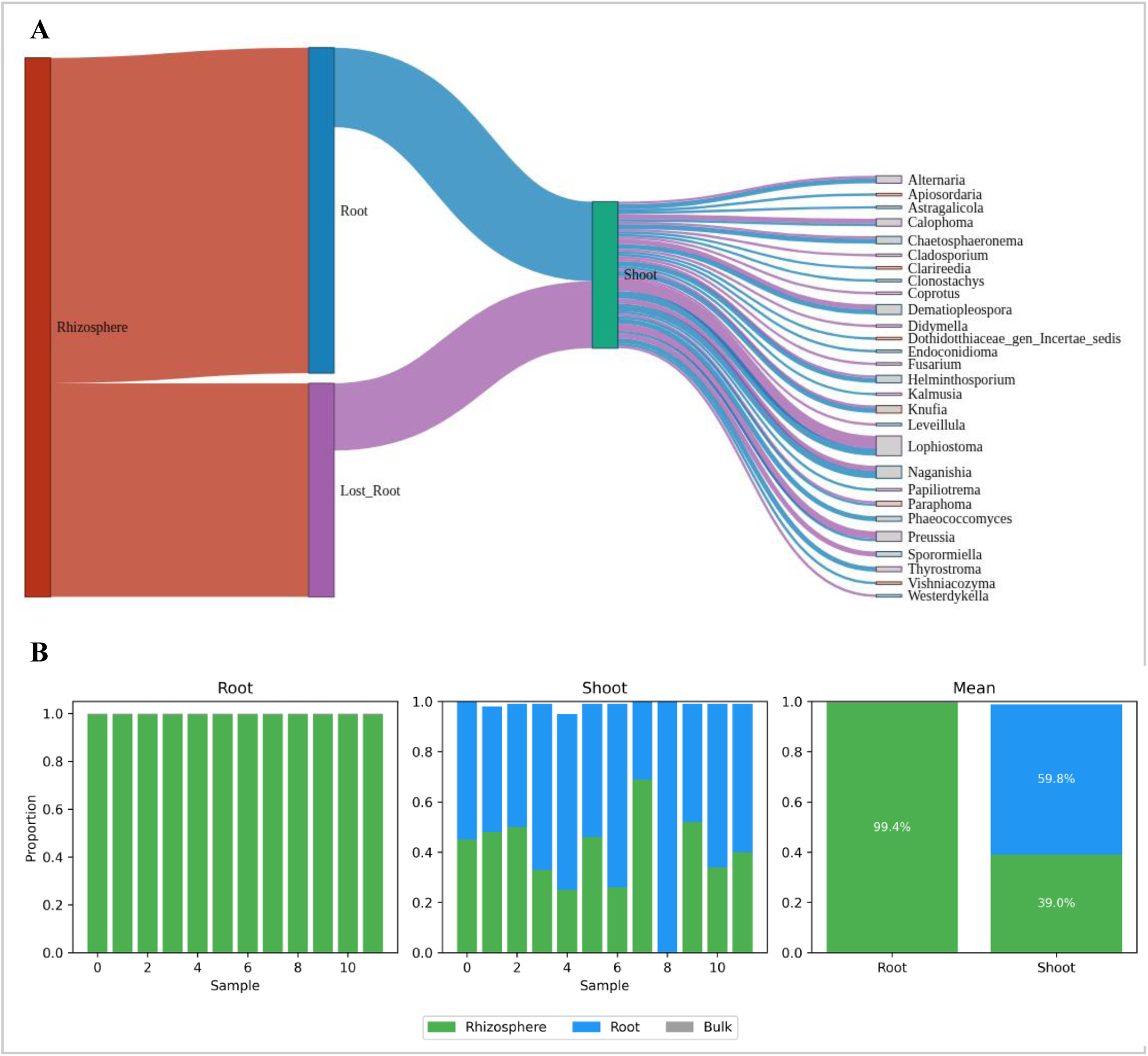
Fungal community assembly across the soil-plant continuum. (A) Sankey diagram showing taxon-specific distribution and transmission across rhizosphere, root, and shoot compartments. (B) Source tracking analysis indicating contributions of rhizosphere and root communities to downstream compartments.

### Culture-dependent characterization and comparison with culture-independent fungal communities

Culture-dependent isolation yielded fungal strains from multiple compartments (rhizosphere, root, and shoot) across the three *Artemisia* species (*A. herba-alba*, *A. mesatlantica*, and *A. negrei*). A total of 21 isolates were obtained, representing multiple fungal taxa. Isolates were identified based on ITS sequencing and confirmed through Sanger sequencing, with sequence similarities ranging from 83.08-99.8% against reference databases (Table S3). The recovered isolates belonged to several genera, including *Alternaria*, *Aspergillus*, *Fusarium*, *Penicillium*, *Preussia*, *Microsphaerop and Botryotrichum*. Phylogenetic analysis further resolved their relationships, showing that isolates clustered into distinct genus-level clades consistent with these taxonomic assignments (Fig. 5A). Multiple isolates were grouped within major genera such as *Aspergillus*, *Penicillium*, and *Fusarium*, forming clusters with corresponding reference sequences.

**Fig. 5.**
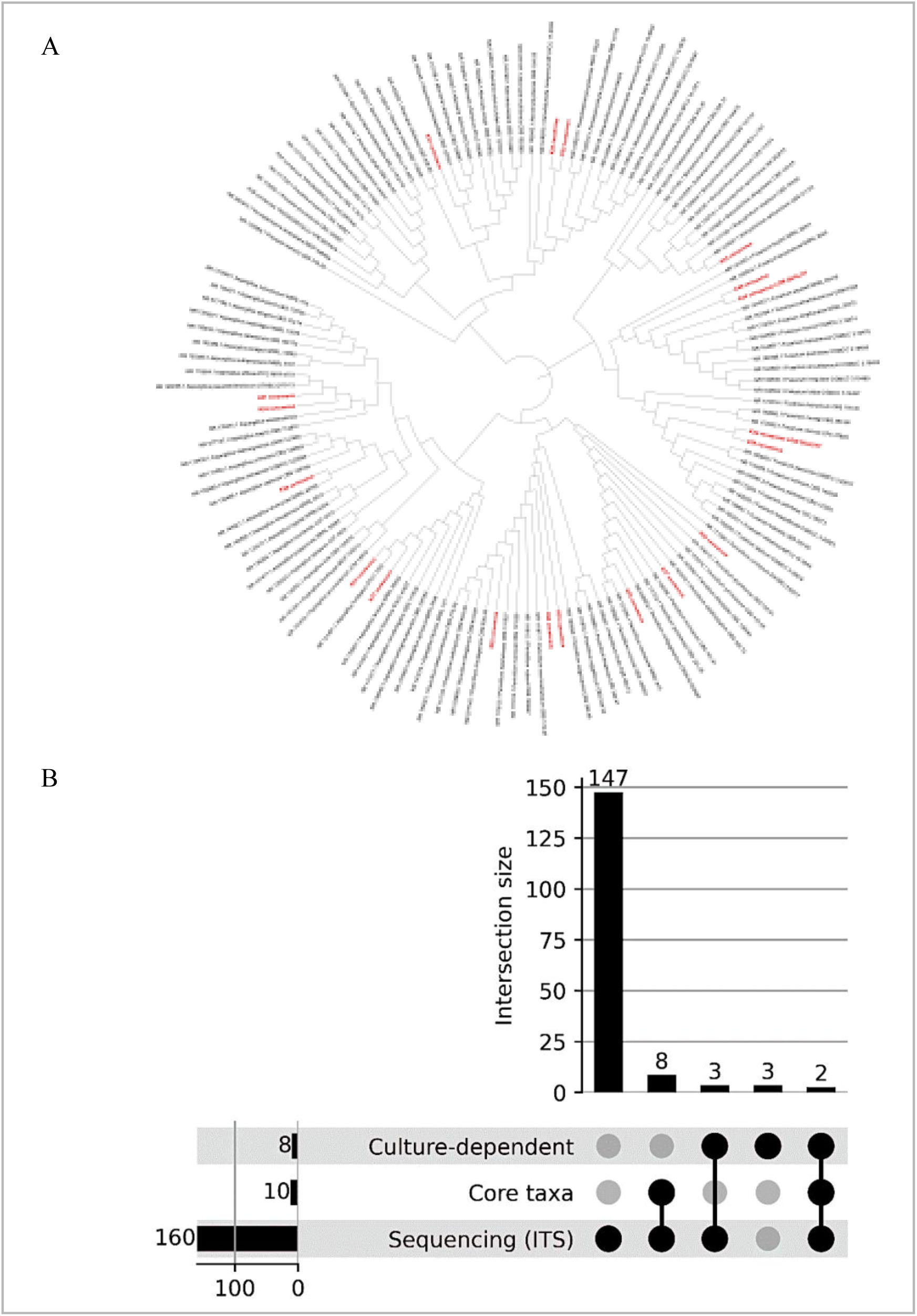
Phylogenetic relationships and overlap of cultured fungal isolates. (A) Phylogenetic tree based on ITS sequences of representative fungal isolates and reference sequences. Isolates cluster within major fungal genera, including *Aspergillus*, *Penicillium*, *Fusarium*, *Alternaria*, and *Preussia*. (B) UpSet plot showing overlap among culture-dependent, culture-independent (ITS), and core fungal taxa. The upper panel represents intersection sizes, and the lower matrix indicates set membership.

Comparison with sequencing-derived data revealed a limited overlap between approaches. While sequencing identified substantially higher diversity (147 genera), only five genera (*Alternaria*, *Aspergillus*, *Fusarium*, *Penicillium*, and *Preussia*) were shared between datasets. Core taxa largely overlapped with sequencing (8 genera), and only two genera were detected across culture-dependent, sequencing-derived, and core datasets (Fig. 5B). These results indicate that culture-dependent methods recover a restricted subset of fungal diversity, primarily representing dominant and ecologically relevant taxa identified by sequencing. Overall, these taxa were distributed across different compartments and host species, with variation in isolate frequency among sample types.

## Discussion

### Compartmental structuring of fungal communities in *Artemisia*

Fungal communities associated with *Artemisia* were structured primarily by plant compartment rather than host species identity. Although alpha diversity did not differ significantly among compartments, clear differences in community composition were observed, indicating that variation among samples was driven by compositional turnover rather than richness. Similar patterns have been reported in plant microbiome studies, where distinct microbial assemblages occur across compartments despite comparable diversity (26, 27). Compartment-specific structuring reflects ecological filtering along the soil–plant continuum, where microbial communities are progressively reshaped during plant colonization (28, 29). In this study, the separation of root and shoot communities from soil-associated communities, together with reduced taxonomic overlap across compartments, supports the presence of such compartment-dependent filtering. The relatively weak differentiation among *Artemisia* species is consistent with previous findings in natural plant systems (30, 31).

### Rhizosphere as a central reservoir and transition zone

The rhizosphere represents a key interface linking soil and plant-associated fungal communities. The consistent occurrence of several taxa across *Artemisia* species, together with the presence of a defined rhizosphere core microbiome, supports its role as a zone of microbial enrichment. Root exudates and plant-derived compounds are known to influence rhizosphere communities by selectively recruiting microorganisms from the surrounding soil (32, 33). The higher overlap observed between bulk soil and rhizosphere communities, compared with plant tissues, is consistent with the rhizosphere acting as a transitional reservoir mediating microbial recruitment (6, 34, 35). The strong overlap between rhizosphere and root core taxa further supports this interpretation, with most root-associated core genera also present in the rhizosphere, indicating that root communities largely represent a filtered subset of the rhizosphere microbiome. In contrast, the limited overlap between bulk soil and plant compartments, together with the absence of core taxa in bulk soil and shoot environments, suggests greater heterogeneity in soil fungal communities and stronger environmental filtering and stochastic colonization in aboveground tissues, consistent with previous studies showing that phyllosphere fungal communities are distinct from belowground communities and are shaped by strong environmental filtering and stochastic colonization (36, 37).

### Selective assembly and differential roles of fungal taxa

Taxonomic patterns along the soil–plant gradient indicate progressive selection of fungal taxa during plant colonization. Root-associated communities were characterized by a narrower set of dominant taxa, whereas shoot communities exhibited distinct composition and increased representation of unclassified fungi. These observations are consistent with hierarchical filtering models, in which microbial communities undergo successive selection steps from soil to rhizosphere and into plant tissues (5, 27, 38). Fungal taxa also differed in their associations with community composition and environmental gradients. Some taxa, such as *Lophiostoma*, were more strongly associated with compositional variation, whereas others, including *Alternaria* and *Didymella*, showed associations with soil-related gradients. These patterns suggest that fungal taxa occupy distinct ecological niches, with some more closely associated with plant compartments and others influenced by environmental conditions (35, 39, 40).

### Sequential assembly and environmental context in arid systems

The combined results support a sequential model of fungal community assembly along the soil-plant continuum. Fungal taxa were progressively filtered from the rhizosphere into root and shoot compartments, with reduced overlap along this gradient. Source tracking analysis further indicated that root communities were largely derived from the rhizosphere, whereas shoot communities showed contributions from both root and rhizosphere sources, suggesting multiple colonization pathways. Soil physicochemical properties were associated with fungal community composition, although their influence varied among taxa. Environmental factors such as nutrient availability, pH, and soil texture are known to influence fungal communities (6, 8), particularly in soil and rhizosphere compartments. However, the stronger association of community composition with plant compartments, together with the lack of significant relationships between soil properties and diversity, suggests that host-associated processes play a major role in structuring fungal communities within plant tissues (5, 39). The SEM results further support that compartment-specific processes exert a stronger influence on community composition than environmental gradients alone. These findings indicate that fungal community assembly in semi-arid Artemisia systems reflects the combined effects of environmental conditions and compartment-specific processes (6, 40). Overall, our results demonstrate that fungal community assembly in *Artemisia* is primarily driven by compartment-specific filtering, with environmental gradients and host species playing secondary roles. These findings support our hypotheses, indicating that fungal communities are structured along the soil-plant continuum, with rhizosphere and root compartments harboring recurrent taxa, and that culture-dependent and sequencing-based approaches capture overlapping but partially distinct fungal assemblages.

## Conclusion

Fungal communities associated with *Artemisia* were structured primarily by plant compartment, with additional contributions from soil physicochemical gradients. Distinct community compositions across rhizosphere, root, and shoot compartments, together with reduced taxonomic overlap along this gradient, support a sequential model of community assembly driven by compartment-specific filtering and compositional turnover along the soil-plant continuum. Several limitations should be noted. Sampling was conducted at a single time point, limiting assessment of temporal variability, and assembly processes were inferred from observational and statistical approaches rather than experimental validation. In addition, the presence of unclassified taxa, particularly in shoot samples, constrained taxonomic resolution and functional interpretation. Future studies should incorporate temporal and experimental approaches to better resolve microbial assembly and transmission. Integrating functional analyses, such as metagenomics or transcriptomics, and expanding sampling across broader environmental gradients and host systems will further clarify the relative roles of environmental and host-associated processes in shaping fungal microbiomes in semi-arid ecosystems.

## Data Availability

Raw ITS sequencing data have been deposited in the NCBI Sequence Read Archive (SRA) under BioProject accession PRJNA1457683 (https://www.ncbi.nlm.nih.gov/bioproject/1457683). Sanger sequences of cultured fungal isolates have been deposited in GenBank under accession numbers PZ575267–PZ575287.

## Declarations

## Ethics approval and consent to participate

This research does not include or report the results of a clinical trial; therefore, trial registration and ethical approval was not required.

## Consent for publication

Not applicable

## Authors contributions

RK: experimentation, data analysis, and draft the manuscript. RE: dada2 and statistical analysis. GA: data analysis. FZA: library preparation and sequencing. MB: sampling and deposition of Vaucher specimens; MH: manuscript review and editing. BA: conceptualization, sampling, manuscript drafting, review and supervision. All authors contributed to the article and approved the submitted version.

## Acknowledgements

We would like to thank the African Genome Center, Mohammed VI Polytechnic University (UM6P), Benguerir, Morocco, for providing access to sequencing facilities and technical support. We also acknowledge Dr. Brahim Ouahzizi and Dr. Hamza Elbony for supporting during sampling.

